# Semi-automated vs. manual: Comparative study of cell culture counting methods using validation parameters

**DOI:** 10.1101/2024.05.30.596619

**Authors:** Selina Ramm, Ulrich Odefey, Björn Frahm, Miriam Pein-Hackelbusch

**Affiliations:** Institute for Life Science Technologies ILT.NRW, OWL University of Applied Sciences and Arts, Lemgo, Germany; Biotechnology and Bioprocess Engineering, OWL University of Applied Sciences and Arts, Lemgo, Germany

## Abstract

Determining cell density and cell viability is fundamental for any cell cultivation process. In addition to the manual counting method using hemocytometers, (semi-)automated methods offer advantages such as lower variability and shortened analysis times. However, these methods should provide at least comparable results to the manual method, which is why a comparison of methods is essential. We conducted a dilution series experimental design according to ISO 20391-2:2019 and compared two cell counting methods based on validation parameters aligned with the ICH Q2(R1) guideline. Regarding specificity and linearity, the manual (hemocytometer) and semi-automated (Countstar BioTech^®^) method exhibited similar results in the two evaluated characteristics total cell density and cell viability of CHO-K1 cells. Regarding repeatability of determining total cell density, the semi-automated method achieved significant (*α* = 0.05) better results with average relative standard deviations of < 6 %, than the manual method with average relative standard deviations of > 9 %. Concerning repeatability of the cell viability measurement, no significant difference between the two methods were shown. These results show the suitabililty of the dilution series experimental design. For the applied example, they indicate that the investigated semi-automated method is an appropriate alternative to the manual method.

## Introduction

Cell counting is fundamental in various industrial and scientific applications. For example, application fields include the development of soft sensors, seed-train planning, optimization of fed-batch control or repeated batch strategies [1] as product formation is often proportional to viable cell density (VCD). In addition to the VCD and total cell density (TCD), cell viability, as the relation of those two parameters, is a key performance indicator of the cell culture’s state. Therefore the reliable quantification is essential for ensuring the success of cell-based processes. To ensure reliability, a procedure for analyzing the suitability of an individual measurement method or for choosing between different measurement methods is necessary. Among the methods available for this purpose, manual cell counting using a hemocytometer stands out as a widely utilized technique [2–5]. Due to several disadvantages such as inter- and intra-analyst variability [6], a trend towards (semi-)automated counting systems can be observed. Compared to the use of manual cell counting methods, (semi-)automated cell counting systems reduce the analytical time and analyst dependent variability due to automation of image processing [6, 7]. Additionally, further insights about the sample can be gained, such as cell diameters, circularity, and cell aggregation [6]. Among the most well-known automated cell counting devices are the Cedex^®^ HiRes Analyzer (F. Hoffmann-La Roche AG, Basel, Switzerland) and the NucleoCounter^®^NC-200^™^(ChemoMetec A/S, Allerod, Denmark). Semi-automated systems include the Countess^™^ System (Invitrogen Corporation, Carlsbad, California) and the Countstar^®^ (Alit Biotech (Shanghai) Co., Ltd., Shanghai, P.R.China). While the categorization in semi-automated and automated systems depends on whether the staining of the sample is done manually or fully automated, the sample analysis is based on digital image analysis in all devices. Cell staining is conducted to distinguish viable from nonviable cells. Due to its simplicity, trypan blue is widely used [8]. The method is based on the principle of cell membrane integrity. Due to their intact membrane viable cells exclude the dye while nonviable cells are stained [9]. Hence, trypan blue exclusion enables immediate identification of viable (unstained) and nonviable (stained) cells.

As already mentioned, determination of cell density and cell viabilty is an important basis for the monitoring and control of cell culture processes. The applied counting method and devices are therefore crucial elements and should be carefully chosen. The examination of new devices is therefore a common task in cell culture process development and evaluation. Prior to application, a method should be proven to work as expected. This means in the assessment of cell density and cell viability, that each new (semi-automated) method should give at least comparable results to those of the manual method. Therefore, the cell counting results obtained from experiments must be evaluated using appropriate parameters. For this purpose, the validation parameters proposed in the International Council for Harmonization (ICH) Q2(R1) guideline [10], such as specificity, linearity, precision, and accuracy, can be utilized as other works have already shown [2, 11, 12].

Our aim was to present a method to evaluate and compare two cell counting methods exemplified by a manual and a semi-automated method. The sample preparation and counting process were compliant to ISO 20391-2:2019 [13] and the comparison was based on validation parameters aligned with the ICH Q2(R1) guideline [10]. We evaluated and assessed the performance of both methods by applying the parameters specificity, linearity, and repeatability for determining the TCD and cell viability of CHO-K1 cells using the trypan blue exclusion assay. First, the TCD was examined to ensure that the basic cell count determination is independent of distinguishing between viable and dead cells. Subsequently, viability was evaluated, which includes the differentiation of the cells. The VCD can be determined using the TCD and viability, so this parameter was not considered separately.

## Materials and methods

### Experimental setup

We conducted a dilution series experimental design according to ISO 20391-2:2019 [13]. The experimental setup was divided into three sets. Set A and B addressed the determination of the TCD while set C was used to determine cell viability. In sets A and C, three dilution fractions were examined each (Table 1). The concentration range of A was oriented at the range of other works [4, 11, 14–16]. The concentration range for B was chosen to cover the range of expected concentrations that occur during cell cultivation in the bioreactor. Due to the broad range, a total of five dilution fractions was considered. Set C had the same TCD in each sample but varying viability. For each set, an equally spaced interval between the dilution fractions of the measured variable was selected. Each set was prepared and analyzed on the same day by the same operator. Table 1 summarizes the described setup.

**Table 1.** Experimental details overview.

| Set | TCD stock cell solution | Quantity DF | Range |
| --- | --- | --- | --- |
| A | $6.6 \times 10^6 \text{ cells mL}^{-1}$ | 3 | $5.0 \times 10^5 - 5.0 \times 10^6 \text{ cells mL}^{-1}$ |
| B | $2.2 \times 10^7 \text{ cells mL}^{-1}$ | 5 | $1.1 \times 10^5 - 2.2 \times 10^7 \text{ cells mL}^{-1}$ |
| C | $1.3 \times 10^6 \text{ cells mL}^{-1}$ | 3 | 36 % – 88 % viability |

TCD, total cell density; DF, dilution fraction.

In general, a set consists of three or five dilution fractions DF (n_DF_=3 or n_DF_=5) and three independent replicate test samples j per DF (n_j_=3). The measurement k of a single test sample is performed in triplicate (n_k_=3 per j) on each of the two devices. The experimental setup is summarized in Fig 1.

**Fig 1.**
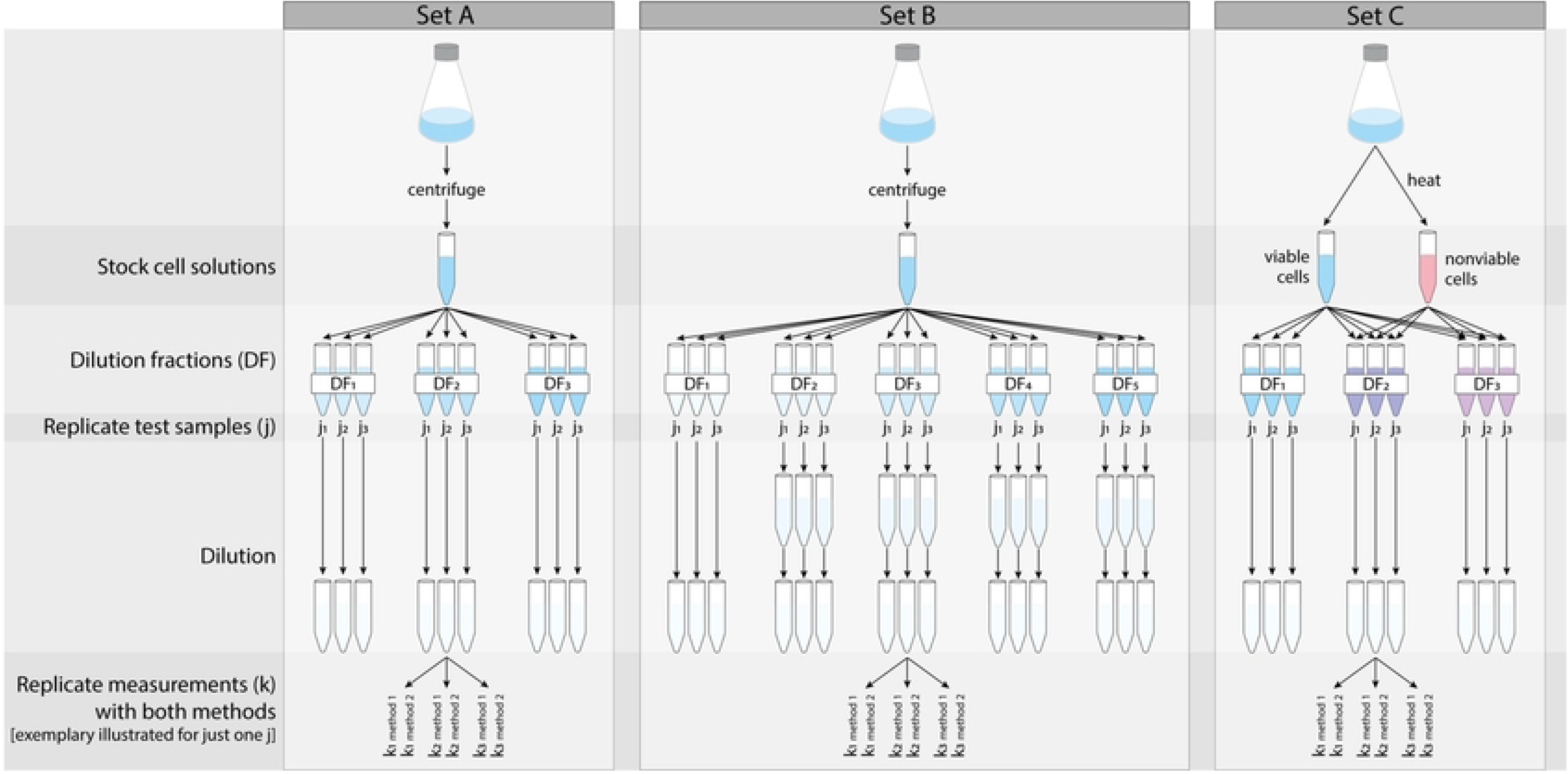
Setup of the dilution series experimental design. Set A and B were used to measure total cell density, set C was used to measure cell viability. Method 1 and 2 refer to the manual and semi-automated method, respectively.

### Preparation of stock cell solutions

The CHO-K1 cells were cultured at 8 – 40 % filling volume in 125 mL shake flasks at 37 °C under 5 % CO_2_ humidified atmosphere (Model CB 150, BINDER GmbH, Tuttlingen, Germany). The shake flasks (Art.-No. 214-0447, VWR International GmbH, Darmstadt, Germany) had no baffles, a vent cap and consisted of polycarbonate. The flasks were incubated and shaken at 70 – 125 rpm (depending on the filling volume) and 25 mm shaking diameter (Model 69455, Infors AG, Bottmingen, Switzerland), to reach a k_L_a of about 17 – 19 h^−1^. The used medium was TC-42 w/o GF (Sartorius Xell GmbH, Schloß Holte-Stukenbrock, Germany) supplemented with 8 mM L-glutamine (Art.-No. 3772, Carl Roth GmbH + Co. KG, Karlsruhe, Germany). To obtain cell concentrations as high as desired for A and B in shake cultures, the shake flask culture was centrifuged for 5 min at 110 x*g* (Model 6K15, Sartorius AG, Göttingen, Germany). The supernatant was discarded and the cell pellet was resuspended in an appropriate volume of medium to generate a stock cell solution with an estimated concentration of 6.6 × 10^6^ cells mL^−1^ for A and of 2.2 × 10^7^ cells mL^−1^ for B. For the viability measurements C, a total cell density (of 1.3 × 10^6^ cells mL^−1^), normally grown in the shake flask, was sufficient. Nonviable cells were obtained by inducing thermal stress in a thermoshaker at 90 °C for 10 min at 500 rpm and 3 mm shaking amplitude (Model MKR 13, HLC BioTech, Bovenden, Germany). Nonviability was confirmed through the staining pattern in a hemocytometer (Neubauer, Paul Marienfeld GmbH & Co. KG, Lauda-Königshofen, Germany). This resulted in a viable stock cell solution with 88 % viability and a nonviable stock cell solution with 0 % viability.

### Preparation of independent replicate test samples

Set A and B: starting from the stock solution, a well-mixed (gently pipetted up and down 10 times) cell suspension was added to a centrifuge tube at the predetermined volume. This volume was verified by weighing using a calibrated scale (Model ATL-224-I, Sartorius AG, Göttingen, Germany). Medium was then added to the cell suspension to reach the targeted dilution fraction (DF_targeted_). The measured dilution fraction for each sample (DF_measured_) was calculated as the mass of the cell suspension over the total sample mass. This procedure complies with the requirements of the ISO 20391-2:2019 [13] for considering pipetting errors in dilution integrity.

In the case of set B, an additional dilution step was introduced to comply with the analytical ranges of the cell counting method to be used. The dilution procedure was the same as for the dilution with medium, except that in this step the samples were diluted with phosphate-buffered saline (PBS) (pH 7.4, Art.-No. E404, VWR Chemicals LLC, Solon, Ohio, USA). The test samples (except the first concentration of 1 × 10^5^ cells mL^−1^) were brought to a cell density of 1 × 10^6^ cells mL^−1^ by this step. The DF_measured_ was calculated as the mass of the test sample mass over the total sample mass.

Set C: a well-mixed (gently pipetted up and down 10 times) viable cell suspension was added to a centrifuge tube at the predetermined volume. This volume was verified by weighing using the calibrated scale. Nonviable cells were then added to the viable cell suspension to reach the DF_targeted_ (here synonymous with targeted viability). The DF_measured_ was calculated as the mass of the viable cell suspension over the total sample mass.

All test samples were prepared in the order of randomly assigned labels. All test samples were diluted with trypan blue (0.4 % in PBS; Art.-No. CN76, Carl Roth GmbH + Co. KG, Karlsruhe, Germany), taking into account the semi-automated device specification, which says that the final concentration of trypan blue in the sample should be 0.1 %. Accordingly, the test samples were diluted 3:1 (300 µL + 100 µL) with trypan blue (0.4 %) in new centrifuge tubes. The preparation of the independent replicate test samples is also summarized in Fig 1.

### Manual cell counting

A Neubauer hemocytometer (0.1 mm depth) (Paul Marienfeld GmbH & Co. KG, Lauda-Königshofen, Germany) was used for manual cell counting. The working range is approximately 10^5^ – 10^6^ cells mL^−1^ [17]. Based on our own experience, the analysis of one chamber, counting four large squares, usually takes about 4 min/100 cells. A volume of 10 µL of dyed cell suspension was loaded into each chamber of the hemocytometer. The chambers were loaded by first pressing on the cover glass and then applying the sample. For each test sample, a total of three chambers (on two hemocytometers) were loaded and counted once by the same operator under a microscope (Model AE31, Moticeurope S.L.U., Barcelona, Spain) with a 10× objective. Once prepared, the samples were immediately imaged and analyzed. For each chamber, all the four large squares having the same size were counted. Cells were counted if they lay within the square or on the left or bottom line edges of the square. The TCD was calculated according to Eq (1), where the volume per large square is 1 × 10^−4^ mL and the dilution factor is ^4^*/*3 for set A and C due to dilution with trypan blue. In set B, in addition to the dilution with trypan blue, the dilution factor depends on the dilution used in the preparation of the test samples. Viable cells were defined as those that were round and clear, nonviable were defined as those that were also round but blue in color.

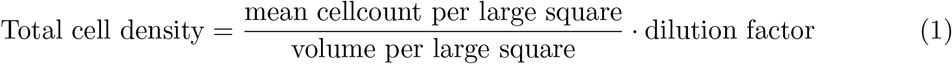

### Semi-automated cell counting

The semi-automated cell count was conducted on a Countstar BioTech^®^ (Alit Biotech (Shanghai) Co., Ltd., Shanghai, P.R.China). According to the user manual of the instrument, the working range is 1 × 10^4^ – 3 × 10^7^ cells mL^−1^. Image acquisition and analysis are typically < 20 sec/sample. The instrument works with a single-use plastic slide with five chambers. For each test sample, three chambers were loaded with 20 µL each. Once prepared, the samples were immediately imaged and analyzed. The parameter settings were set according to the manufacturer’s recommendations:

“Min_Size = 7”; “Max_Size = 20”; “Prm_Grdthres = 7”; “Alive_Dead_Param = 2”; “State_Param = 0”; “Sep_Param = 0.7”; “Overlap_Param = 0.5”;

“Big_block_flag = 1000”; “Eject_dis = 3”; “Reg_in_param = 0.5”. TCD and cell viability were directly output by the Countstar^®^software.

### Statistical analysis

Assay characteristics evaluated in this study included specificity, linearity and precision (repeatability). For the definitions of these validation parameters, refer to the ICH Q2(R1) guideline [10]. Specificity was evaluated on the matrix, here medium diluted 3:1 with trypan blue (0.4 % in PBS). In accordance with the recommendations of the ICH Q2(R1) guideline [10], linearity was evaluated based on a straight-line fit, where a weighted least squares (WLS) approach was used. This means both, TCD and cell viability, were assumed to depend linearly on the targeted dilution fraction. Due to this assumption and the fact that relatively few data points are available, the residual plots were not considered. As required by the guideline, the correlation coefficient (*R*^2^), y-intercept and slope of the straight-line fit were determined for each counting method. The residual sum of squares (RSS) was not reported as the requirement for specifying the RSS is omitted in the revised guideline ICH Q2(R2) [18]. Repeatability was evaluated as the relative standard deviation (RSD) for each method using triplicates of measurements (n_k_=3). The nonparametric Mann-Whitney U-Test was used to compare the medians of the average RSDs over all DF of both methods. A one-sided test was used, i.e. our null hypothesis was that manual cell counting gives a smaller RSD value. The significance level was set to *α* = 0.05. Python programming language (version 3.12) and the modules NumPy (version 1.26.2), SciPy (version 1.11.4), Matplotlib (version 3.8.2), and Pandas (version 2.1.4) were used for the statistical analyses.

## Results and Discussion

The manual (hemocytometer) and semi-automated method (Countstar BioTech^®^) were compared in terms of determination of TCD and cell viability, as both parameters are of great importance for monitoring, controlling, and optimizing cultivation processes. First, it has been investigated, whether both methods yield comparable results for the TCD. Once this was proven, a comparison of methods for cell viability determination was conducted. All raw data of the determination of TCD and viability can be found in S1 Table.

### Comparability in terms of specificity

Since the specificity measurements were evaluated on a cell-free matrix, no cells were expected to be detected. The manual and semi-automated method provided results of 0 cells mL^−1^ and 0 % viability each, indicating that both methods are specific and hence comparable.

### Comparability in terms of linearity

#### Results related to total cell density

As no major deviations of the DF_measured_ from the DF_targeted_ could be measured (S1 Table), for easier interpretability of the results, the DF_targeted_ will be considered in the analysis. The TCD was measured in million cells/mL. The measured TCDs of set A were plotted against the DF_targeted_ for both methods in Fig 2 and a straight-line fit was applied. At this point, we would like to emphasize that the plotted measured TCDs do not represent the raw TCDs measured in reality, as they have been extrapolated using the dilution factors, as is standard practice in other works as well [5, 11, 16, 19]. The intercept, slope and *R*^2^ of the straight-line fit of both methods are reported in Table 2. The intercepts are close to zero, as expected. As the accuracy was not examined due to a lack of reference material, however, no further interpretation of the intercept and the slope can be made. Although there are beads that can be used as reference standards for determining accuracy, there is debate about the extent to which they are truly suitable as reference materials for viable mammalian cells [14]. Still, both fits show a *R*^2^ > 0.9, indicating that the data of both methods are comparably well explained by a straight line.

**Table 2.** Parameter of the straight-line fit.

|  |  | **b (± SE)** | **m (± SE)** | **$R^2$** |
| --- | --- | --- | --- | --- |
| **Set A** | **manual** | $0.034 \pm 0.079$ | $6.030 \pm 0.408$ | 0.995 |
| | **semi-automated** | $0.036 \pm 0.005$ | $7.513 \pm 0.062$ | 1.000 |
| **Set B** | **manual** | $0.058 \pm 0.017$ | $14.313 \pm 0.858$ | 0.963 |
| | **semi-automated** | $0.046 \pm 0.001$ | $20.217 \pm 0.337$ | 0.970 |
| **Set C** | **manual** | $-3.535 \pm 4.298$ | $93.321 \pm 5.261$ | 1.000 |
| | **semi-automated** | $-1.446 \pm 10.964$ | $83.773 \pm 18.146$ | 1.000 |

b, y-intercept; m, slope; SE, standard error; $R^2$ , correlation coefficient.

**Fig 2.**
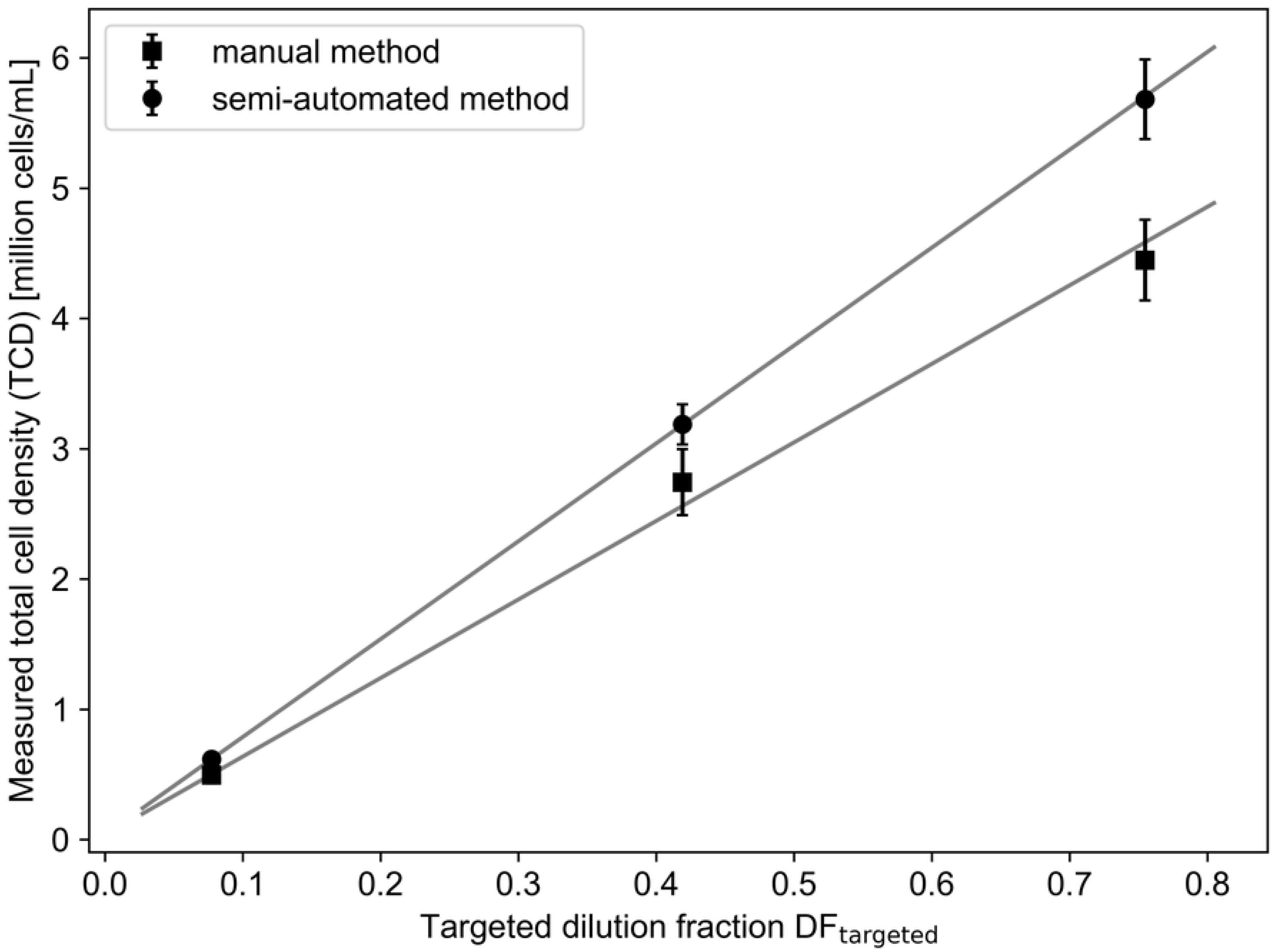
Linearity of total cell density over three-point concentration range. A straight-line fit reasonably captures the dependence of total cell density as a function of dilution fractions. Error bars indicate one standard error of the mean.

As linear trends based on only three concentrations should not be overestimated, we investigated a broader concentration range with a total of five concentrations afterwards in set B (Fig 3). The straight-line fit of the measured TCDs over all DF_targeted_ shows an *R*^2^ of 0.963 for the manual method and an *R*^2^ of 0.970 for the semi-automated method (Table 2). This suggests that the data from both counting methods are equally well described by the straight-line fit.

**Fig 3.**
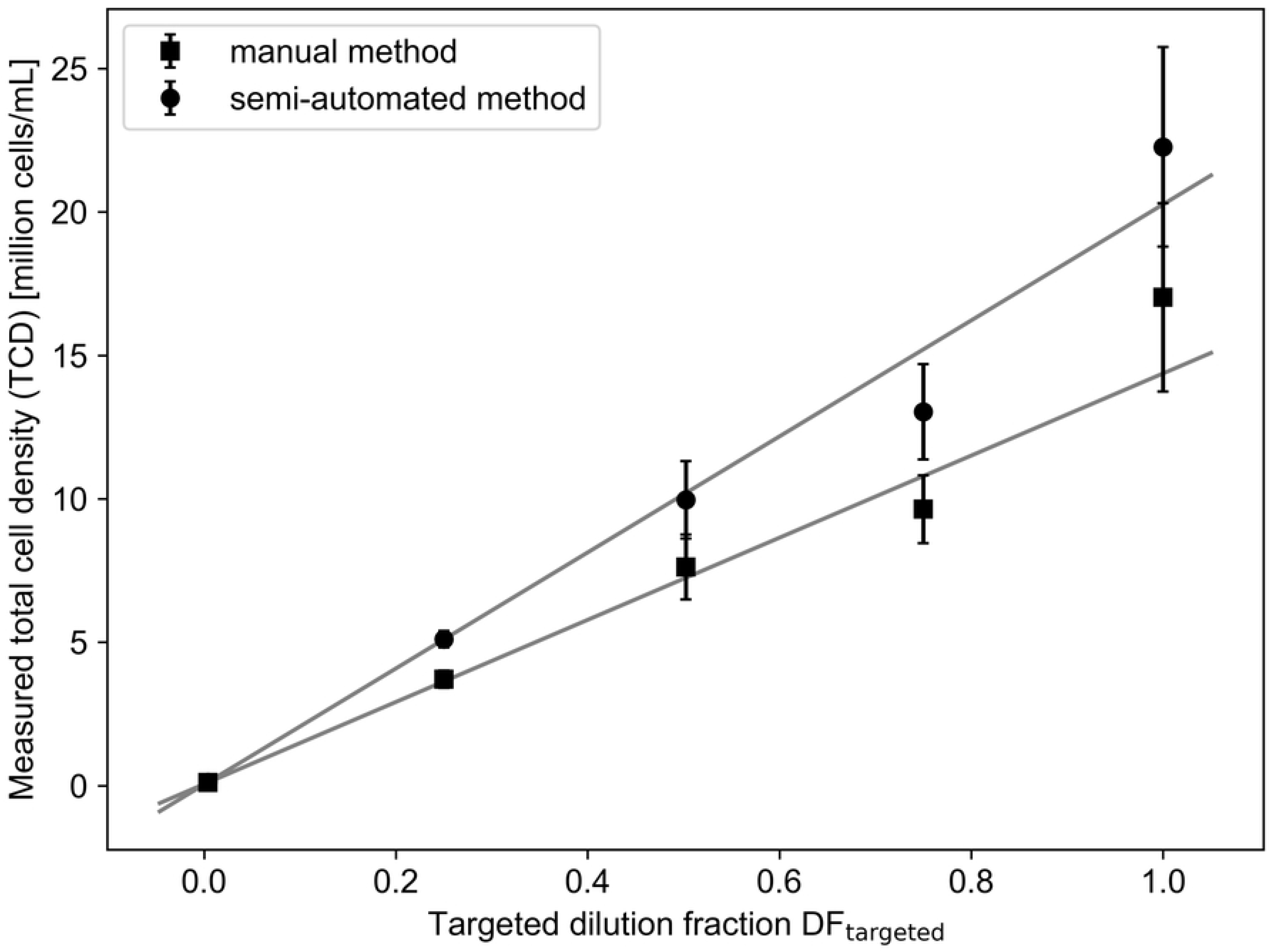
Linearity of total cell density over five-point concentration range. A straight-line fit reasonably captures the dependence of total cell density as a function of dilution fractions. Error bars indicate one standard error of the mean.

#### Results related to viability

Fig. 4 shows cell viability as a function of DF_targeted_. Both fits show an *R*^2^ of 1.000 (Table 2), indicating that the data are well explained by the straight-line fit.

**Fig 4.**
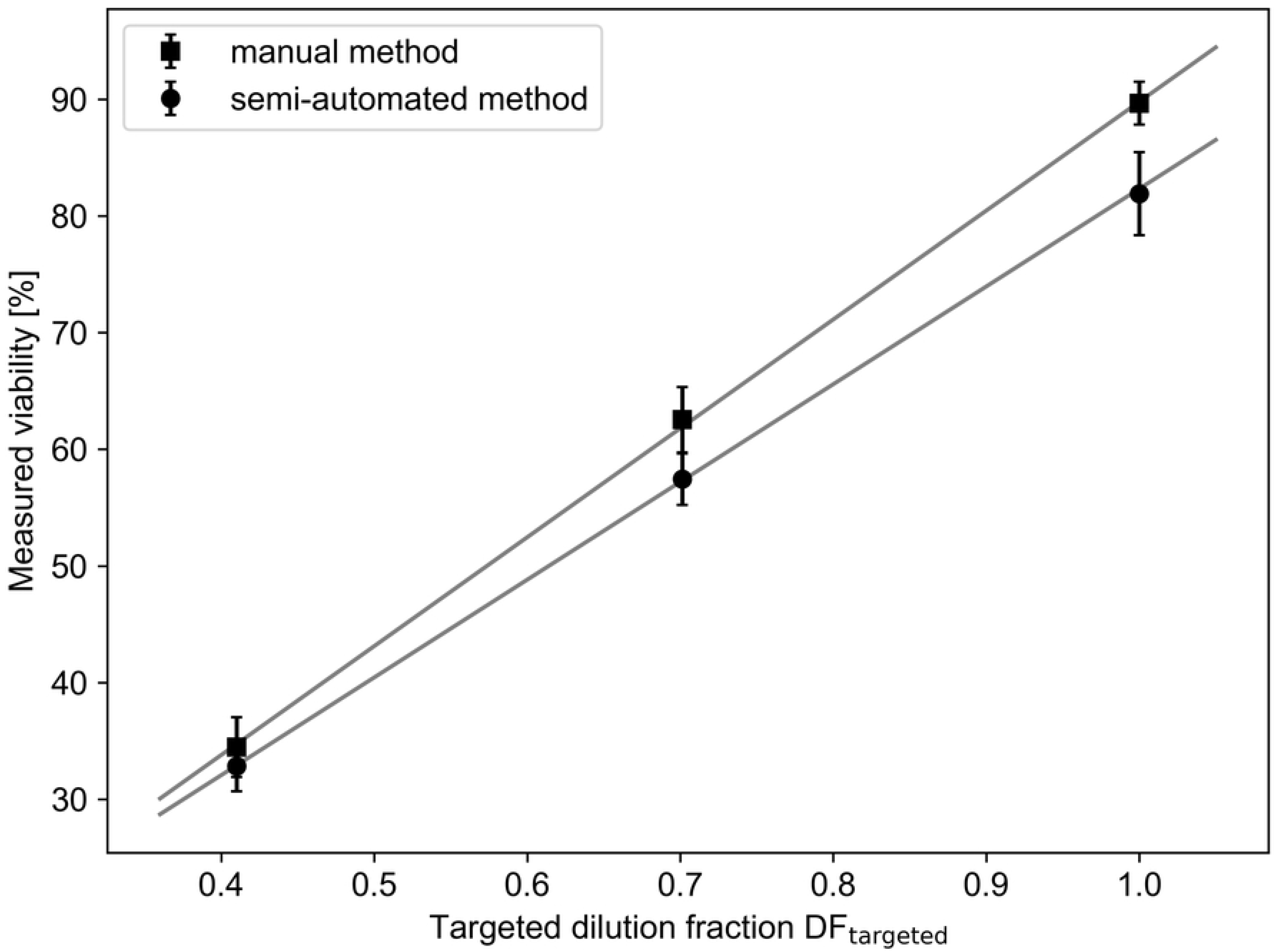
Linearity of viability over three-point concentration range. A straight-line fit reasonably captures the dependence of cell viability as a function of dilution fractions. Error bars indicate one standard error of the mean.

### Comparability in terms of repeatability

#### Results related to total cell density

The average RSD over all DF (n=9) for set A (Sheet A in S2 Table), indicates that the semi-automated method tends to exhibit lower variation (4.9 % ± 1.3 %) than manual cell counting (9.1 % ± 1.1 %). The results are given as mean ± standard error of the mean (SEM). A statistical significant difference between the average RSDs of both methods was confirmed through a Mann-Whitney U-Test (*p* = 0.004). The Mann-Whitney U-Test was used because it is not dependent on a particular distribution [20, 21]. For set B, the average RSD over all DF (n=15) (Sheet B in S2 Table) also indicates a lower variation of the semi-automated method (5.8 % ± 0.8 %) compared to the manual method (10.9 % ± 1.2 %). The Mann-Whitney U-Test gave a p-value of 0.021, confirming statistical significant difference between the average RSDs of both methods. These RSDs can be compared with those of Sarkar et al. [4].

Although human mesenchymal stromal cells were examined there instead of CHO cells, an average RSD of 11.2 % was determined for the manual method (hemocytometer) and an average RSD of 6.6 % for the automated method (NucleoCounter^®^ NC-200) over a range of 0.22 × 10^6^ – 1.98 × 10^6^ cells mL^−1^. If one wanted to validate the methods regarding repeatability, one could choose an acceptance criterion of RSD < 10 % for the TCD determination, in accordance with [12] and [22].

#### Results related to viability

As already observed in the results of the TCD determination, the manual method also demonstrates marginally higher variation (4.6 % ± 1.2 %) in terms of the average RSD over all DF (n=9) for viability determination compared to the semi-automated method (4.4 % ± 1.0 %) (Sheet C in S2 Table). Here, the difference between the average RSDs of both methods is not statistically significant according the Mann-Whitney U-Test *p* = 0.604.

### Comparability in terms of accuracy

Accuracy was not investigated in this study due to the lack of reference materials. Without reference material, no clear statement can be made about which of these two methods is more accurate. A clue, however, is provided by the statement of the ICH Q2(R1) guideline, which suggests that “accuracy may be inferred once precision, linearity and specificity have been established” [10]. Although the methods were not validated with respect to these parameters, but only compared based on them, the lower average RSD over all DF of the semi-automated method suggest a higher repeatability in the determination of TCD and cell viability compared to manual cell counting. As the numerical results for specificity and linearity almost coincide (Table 3), the marginally improved repeatability of the semi-automated method may also indicate a higher accuracy.

**Table 3.** Summarized results of the method comparison.

|  |  | Specificity | Linearity |  |  | Repeatability |
| --- | --- | --- | --- | --- | --- | --- |
|  |  |  | b (± SE) | m (± SE) | R<sup>2</sup> | average RSD (± SEM) |
| Set A | manual | 0 cells mL<sup>-1</sup> | 0.034 ± 0.079 | 6.030 ± 0.408 | 0.995 | 9.1 % ± 1.1 % |
|  | semi-automated |  | 0.036 ± 0.005 | 7.513 ± 0.062 | 1.000 | 4.9 % ± 1.3 % |
| Set B | manual |  | 0.058 ± 0.017 | 14.313 ± 0.858 | 0.963 | 10.9 % ± 1.2 % |
|  | semi-automated |  | 0.046 ± 0.001 | 20.217 ± 0.337 | 0.970 | 5.8 % ± 0.8 % |
| Set C | manual | 0 % viability | -3.535 ± 4.298 | 93.321 ± 5.261 | 1.000 | 4.6 % ± 1.2 % |
|  | semi-automated |  | -1.446 ± 10.964 | 83.773 ± 18.146 | 1.000 | 4.4 % ± 1.0 % |

b, y-intercept; m, slope; SE, standard error; R<sup>2</sup>, correlation coefficient; RSD, relative standard deviation; SEM, standard error of the mean

## Conclusion

Cell concentration as well as cell viability are key parameters for fields such as process monitoring, control and design, seed trains or development of process measurement technology. For this, evaluating cell counting devices is a necessary step. In this paper, the comparison of two cell counting methods based on validation parameters aligned with the ICH Q2(R1) guideline has been presented, exemplified by a manual (hemocytometer) and a semi-automated (Countstar BioTech^®^) method. Overall, both methods exhibited similar results in the two evaluated characteristics TCD and cell viability, regarding specificity, linearity and repeatability. However, the semi-automated method appears to achieve slightly better results in repeatability, as shown by lower average RSD over all DF (Table 3). As no accuracy assessment was performed in this study due to the lack of reference materials, no valid statement can be made about this parameter. Nonetheless, the improved repeatability of the semi-automated method could indicate that it provides results closer to the true value. In summary, it can be stated that the semi-automated method is an appropriate alternative to the manual method for determining the TCD and viability of CHO-K1 cells using the trypan blue exclusion assay. In addition to comparable results, this method offers advantages over the manual method, as shortened analysis times (by at least 12 times) and the acquisition of additional insights about the sample, such as cell diameter.

## Supporting information

**S1 Table. Raw data**. Measured data of the determination of total cell density (TCD) and cell viability by the manual and semi-automated method. (A) TCD over three-point concentration range. (B) TCD over five-point concentration range. (C) Viability over three-point concentration range.

**S2 Table. Descriptive statistics**. Mean, standard deviation and relative standard deviation of the triplicate measurements of total cell density (TCD) and cell viability by the manual and semi-automated method. (A) TCD over three-point concentration range. (B) TCD over five-point concentration range. (C) Viability over three-point concentration range.

## Acknowledgments

The authors gratefully acknowledge discussions with Dr. Tanja Hernández Rodríguez, Julius Wörner, Bodo Kohring and Veronika Gassenmeier.

